# Targeted RNAi screen reveals novel regulators of RNA-binding protein phase transitions in *Caenorhabditis elegans* oocytes

**DOI:** 10.1101/2025.09.08.674920

**Authors:** Mohamed T. Elaswad, Grace M. Thomas, Corrin Hays, Nicholas J. Trombley, Jennifer A. Schisa

## Abstract

The ability of oocytes to maintain their quality is essential for successful reproduction. One critical aspect of oocyte quality and successful embryogenesis after fertilization is the proper regulation of the stores of maternal mRNA by RNA-binding proteins. Many RNA-binding proteins undergo regulated phase transitions during oogenesis, and alterations of the protein phase can disrupt its ability to regulate mRNA stability and translation. In *C. elegans*, genetic screens have identified regulators of RNA-binding protein condensation in arrested oocytes of females and in embryos, but less attention has focused on phase transitions in maturing oocytes of young adult hermaphrodites. Interestingly, of the relatively few regulators of RNA-binding protein phase transitions identified to date in maturing oocytes, several genes overlap with those required for clearance of protein aggregates in maturing oocytes. To determine the extent to which the temporally linked processes of clearance of damaged proteins and maintenance of RNP complexes are coordinated at a molecular level, we conducted a targeted RNAi screen of genes required for removal of protein aggregates in maturing oocytes. We identified six novel regulators of phase transitions of the KH-domain protein MEX-3 and obtained strong evidence that the regulatory network of protein aggregate clearance overlaps with, but is distinct from, the regulation of MEX-3 phase transitions in the oocyte.

## INTRODUCTION

The regulation of maternal mRNAs by RNA-binding proteins is essential for oocyte growth and early embryogenesis across metazoa (reviewed in Conti and Kunitomi 2024). In the absence of tight spatial and temporal regulation of mRNA stability and translation, birth defects and infertility can arise. One important consideration regarding mRNA regulation across cell types is the emerging paradigm of cellular organization of RNA and RNA-binding proteins via dynamic, membraneless organelles (Holehouse and Alberti 2025). Examples of membraneless organelles include stress granules and Processing bodies which are detected in a variety of somatic cells in addition to germ cells. During oogenesis, many RNA-binding proteins and maternal mRNAs undergo dynamic phase transitions, alternating between decondensed and condensed states throughout the meiotic cell cycle and in response to environmental or physiological cues (Schisa et al. 2001; Jud et al. 2008; Flemr et al. 2010; Cheng et al. 2022). Moreover, recent studies have illuminated the significance of protein phase for RNA-binding protein function and regulation of maternal mRNAs. In *Drosophila* oogenesis, perturbing the solid state of *oskar* RNP granules impairs *oskar* mRNA localization and translation, leading to embryo defects (Bose et al. 2022). In mammalian oocytes, disruption of a membraneless organelle named mitochondria-associated ribonucleoprotein domain (MARDO) in GV-stage oocytes results in premature degradation of maternal mRNAs prior to maturation (Cheng et al. 2022). Two of the RNA-binding proteins that undergo phase separation to comprise the MARDO, DDX6 and LSM14, also undergo phase transitions in oocytes of other vertebrates and invertebrates (Ladomery et al. 1997; Nakamura et al. 2001; Navarro et al. 2001; Audhya et al. 2005; Boag et al. 2005; Squirrell et al. 2006; Jud et al. 2008; Noble et al. 2008; Hubstenberger et al. 2013; Elaswad, Watkins, et al. 2022).

The *C. elegans* model offers many advantages in studying post-transcriptional gene regulation, and genetic screens have identified regulators of RNA-binding protein phase transitions in meiotically-arrested oocytes of *C. elegans* females (Hubstenberger et al. 2015; Wood et al. 2016). However, fewer studies have focused on regulators required to promote the decondensed phase of RNA-binding proteins in oocytes undergoing active maturation in hermaphrodites. In maturing oocytes of young hermaphrodites, untranslated maternal mRNAs and the KH-domain protein MEX-3 are relatively decondensed. However, when aged hermaphrodites become depleted of sperm, the mRNAs and MEX-3 undergo phase separation, forming large condensates in arrested oocytes (Figure 1A; (Schisa et al. 2001; Jud et al. 2008). Understanding the regulation of MEX-3 phase transitions is of high interest because two human Mex3 orthologs colocalize with P-bodies, and in colorectal cancer cells, unregulated hMex3 phase transitions lead to mRNA degradation and poor patient outcomes (Buchet-Poyau et al. 2007; Chen et al. 2024). In *C. elegans* oocytes, MEX-3 physically interacts with two proteins, LIN-41 and OMA-1, that repress translation of maternal mRNAs (Spike et al. 2014; Tsukamoto et al. 2017). Our past studies demonstrate that MEX-3 phase transitions are reversible and correlated with activated Extracellular signal-Regulated Kinase (ERK) and Major Sperm Protein signals from sperm (Jud et al. 2008). The phase transitions of MEX-3 also correlate with alterations in ER morphology, and the RNA helicase CGH-1/DDX6, the CCT chaperonin, and actin are required to inhibit ectopic MEX-3 condensation and ectopic ER sheets (Patterson et al. 2011; Langerak et al. 2019; Elaswad et al. 2024).

**Figure 1.**
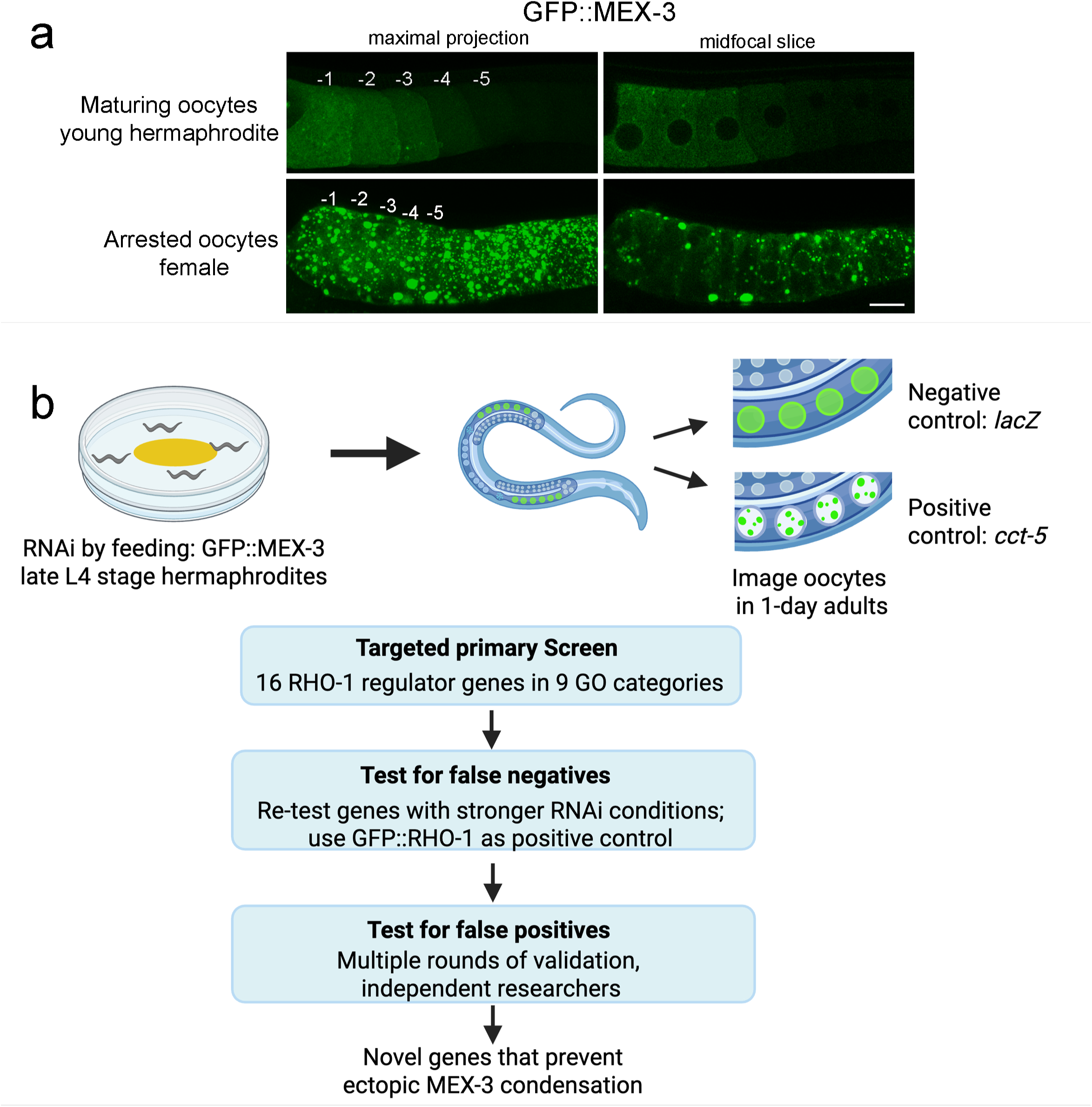
Rationale and workflow for targeted RNAi screen to identify regulators of MEX-3 phase transitions in oocytes. A) The KH-domain protein MEX-3 is dynamic; while it is mostly decondensed in the maturing oocytes of young hermaphrodites, MEX-3 undergoes dramatic condensation into large granules of oocytes undergoing an extended meiotic arrest, e.g. in females lacking sperm. The most proximal *C. elegans* oocyte, closest to the sperm, is referred to as the −1 oocyte. The extent of arrest can be seen when comparing the positions of the five most proximal, maturing and arrested oocytes. A maximal confocal projection and single z-slice are shown for comparison. Scale bar is 10 µm. B) The targeted RNAi screen used a GFP::MEX-3 strain (DG4269) where L4-stage hermaphrodites were moved onto plates with RNAi bacteria. After 24 hours, oocytes in 1-day adults were imaged using confocal microscopy. The negative control was *lacZ*, and the positive control was *cct-5.* In the primary screen, sixteen RHO-1 regulator genes were depleted, one at a time. To test for false negatives, negative genes were r-screened using stronger RNAi conditions, and depletions were performed in parallel in GFP::RHO-1 to ensure efficacy of gene expression depletion. Positive hits were re-screened in at least 3 rounds of validation by multiple researchers. The screen uncovered novel genes that prevent ectopic condensation of MEX-3. Created in BioRender. Schisa, J. (2025) https://BioRender.com/an863wa

Interestingly, studies have identified similar correlations between sperm signals, activated ERK, and the clearance of protein aggregates in oocytes. Moreover, CGH-1, the CCT chaperonin, and actin prevent both RHO-1 protein aggregates and ectopic MEX-3 condensates (Bohnert and Kenyon 2017; Samaddar et al. 2021). These shared characteristics prompted the current investigation to determine the extent to which the regulatory networks of MEX-3 phase transitions and protein aggregates overlap. Genes in eleven gene ontology (GO) categories prevent ectopic aggregates of GFP::RHO-1 in maturing oocytes (Samaddar et al. 2021). Since our prior results identified multiple genes in the protein folding (chaperone) and cytoskeleton GO categories as regulators of MEX-3 phase transitions, here we focused on testing genes from the other nine categories for roles in inhibiting ectopic MEX-3 condensation. We identify *copb-2*, *sar-1, sec-24.1, let-711, pbs-7,* and *rpn-6.1* as novel regulators to prevent ectopic condensation of MEX-3 in maturing oocytes. Our results also strongly suggest that genes involved in lysosome acidification, regulation of mitochondrial membrane potential, calcium ion transport, and ESCRT complex-mediated autophagy are not required to modulate MEX-3 phase transitions. Taken together, our results demonstrate that the regulatory pathway of MEX-3 phase transitions is distinct from RHO-1 aggregates during oocyte maturation.

## METHODS

### Worm Strains and Maintenance

*Caenorhabditis elegans* strains were maintained on Nematode Growth Medium (NGM) plates and fed OP50 *Escherichia coli* at 20°C, unless specified (Brenner 1974). The following strains were used: DG4269 (*tn1753[gfp::3xflag::mex3]*) and SA115 (*unc-119(ed3) III; tjIs1[pie-1::GFP::rho-1 + unc-119(+)]*). Strains were synchronized before experiments using a hypochlorite bleaching method.

### RNA-mediated interference (RNAi) Screen Design

Feeding RNAi was performed as described previously (Timmons and Fire 1998). Transformed RNAi clones of HT115 *E. coli* expressing dsRNA for the target gene from the Source Bioscience RNAi library (Kamath and Ahringer 2003) were used to deplete gene expression by feeding. Each colony of bacteria was grown in LB with 50 μg/mL carbenicillin at 37°C for 7 hours. To test for false negatives using more stringent conditions, cultures were supplemented with 5 mM IPTG (isopropyl β-D-1-thiogalactopyranoside) for an additional ∼ 60 minutes at 37°C. Plates with RNAi media (NGM containing 50 μg/ml carbenicillin and 1 mM IPTG) were seeded with the liquid culture. dsRNA expression was induced at room temperature overnight, and the plates were blinded. All RNAi clones were verified by sequencing.

In the primary screen we tested 16 genes in nine of the GO categories identified as RHO-1 regulators (Figure 1, Table 1). No supplemental IPTG was included in RNAi subcultures of the primary screen. Late L4 (fourth larval) - stage GFP::MEX-3 hermaphrodites were placed on RNAi media at 24°C for 24 hours before collecting confocal images of oocytes in 1-day post-L4 adults. The negative control was *lacZ(RNAi)* where MEX-3 is largely diffuse in oocytes. The positive control was *cct-5(RNAi)* where MEX-3 ectopically condenses (Elaswad et al. 2024). Candidates for positive hits were defined as gene depletions yielding ectopically condensed MEX-3 in oocytes of at least half of the five 1-day post-L4 hermaphrodites scored. We noted the phenotypes of ectopic MEX-3 condensates included small, spherical granules with, or without, accompanying strong enrichment of MEX-3 at the membrane.

**Table 1.**
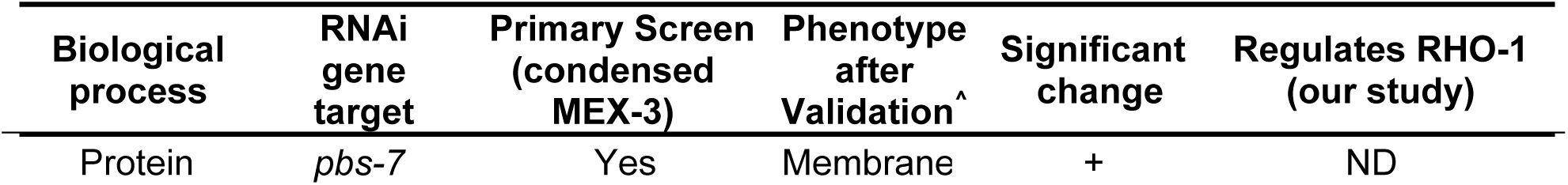
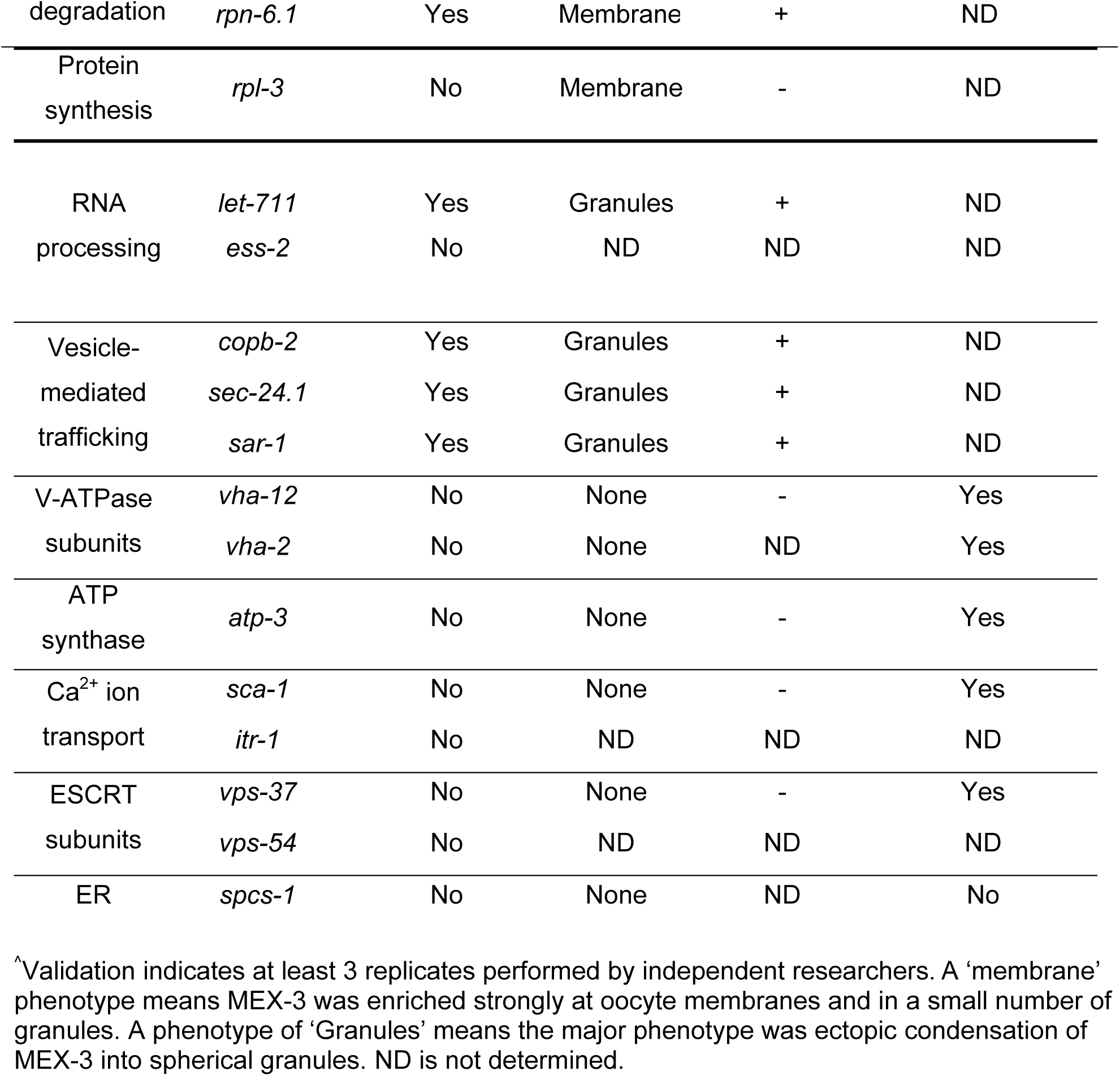
RNAi screen identifies novel regulators of MEX-3 phase transitions and distinguishes MEX-3 regulation from the network regulating clearance of RHO-1 protein aggregates.

To minimize missing false negatives, we re-tested the negative genes using stronger RNAi conditions, bacterial cultures supplemented with 10 mM IPTG for 45-60 min. prior to plating on RNAi media. As a readout of the RNAi efficacy in depleting expression of the intended target genes, we tested each depletion with GFP::RHO-1 worms in parallel to testing GFP::MEX-3 worms. Depletions of five genes of six tested resulted in the expected phenotype of ectopic RHO-1 aggregates (Samaddar et al. 2021). For one gene, *spcs-1*, we did not reproduce the ectopic RHO-1 phenotype across three replicates. To test for false positives and validate candidate positive hits, at least two independent researchers tested positive hits in at least three replicates.

### Microscopy and Image analysis

Confocal Z-stacks of oocytes were collected on a Nikon A1R confocal microscope system. Worms were picked onto slides made with 2% agarose pads and paralyzed using 6.25mM levamisole. All images were collected using identical levels and settings. Worms on each slide were imaged within 10 minutes of mounting on slides to avoid inadvertent stress responses (Elaswad, Munderloh, et al. 2022).

### Phenotype Scoring

After the validation step, we used both qualitative and quantitative methods to score MEX-3 phenotypes. For gene depletions where we did not detect at least a 50% penetrant condensation phenotype, we noted ‘none’ (Table 1). For depletions where many spherical granules were detected, we noted ‘Granules’, and for depletions where MEX-3 was strongly enriched at the cell membrane and in a small number of granules, we noted ‘Membrane’ (Table 1). To determine if the distribution of MEX-3 became significantly more condensed than in the negative control, we used the skewness analysis tool in Fiji (ImageJ). We drew an ROI around each of the three most proximal oocytes, excluding the nucleus. Skewness is a measure of the uniformity of fluorescence signal in an ROI. A value of zero indicates complete symmetry of fluorescence, i.e. no significant condensation. Skewness values deviating from zero indicate increasingly asymmetric distributions of fluorescence, i.e. a proxy for condensation (Elaswad et al. 2024). The mean skewness value for three oocytes was calculated for each worm. Statistical analyses are described below. Significant MEX-3 condensation was detected for six of seven depletions.

### Statistical analysis

Sample sizes were determined using G*Power 3.1 for power analyses; all experiments were blinded and done at least in triplicate. Data are presented as mean ± SEM unless otherwise indicated. Statistical analyses were performed on GraphPad Prism 10.5, and specific tests are noted in figure legends. The Dunn’s correction was used with Kruskal-Wallis tests to control the type I error rate. Corrected P-values are presented, and P-values <0.05 were considered statistically significant.

### Data Availability

All *C. elegans* strains are available at the CGC and upon request. The authors state that all data necessary for confirming the conclusions are present within the article, figures, tables, including supplementary information.

## RESULTS AND DISCUSSION

### Design of targeted RNAi screen to identify regulators of MEX-3 phase transitions in maturing oocytes

To determine the extent to which the genes required to prevent RHO-1 aggregates in maturing oocytes also prevent ectopic condensation of RNA-binding proteins, we performed a targeted RNAi screen. We selected sixteen candidate genes across nine GO categories/biological processes that are implicated in the model for preventing protein aggregates in maturing oocytes (Samaddar et al. 2021)(Table 1). We chose to investigate regulation of the KH-domain RNA-binding protein MEX-3 in our screen as it is evolutionarily conserved, with four homologs in mammals, and dynamic MEX-3 phase transitions have been characterized *in vitro* and across species *in vivo* (Donnini et al. 2004; Buchet-Poyau et al. 2007; Jud et al. 2008; Wood et al. 2016; Elaswad et al. 2024). In maturing oocytes of young hermaphroditic worms, the MEX-3 protein is largely decondensed throughout the cytosol of the most proximal oocytes; note, the most proximal oocyte positioned for meiotic maturation is referred to as the −1 oocyte (Figure 1A) (Draper et al. 1996). In aged hermaphrodites depleted of sperm or females lacking sperm, MEX-3 is highly condensed into granules in the meiotically arrested oocytes (Figure 1A) (Jud et al. 2007). If the female mates with a male, and meiotic maturation resumes, MEX-3 decondenses (Jud et al. 2008). In our screen the negative control was *lacZ(RNAi)*, and the positive control was *cct-5(RNAi)* (Elaswad et al. 2024) (Figure 1B). A positive hit was broadly defined as ectopic condensation of MEX-3 in the maturing oocytes of at least 50% of worms. The primary screen was followed by tests for false negatives and replicates to identify and remove false positives (Figure 1B).

After *lacZ(RNAi)* MEX-3 was decondensed in oocytes, and in the positive control, *cct-5(RNAi),* two types of ectopic condensates were detected: small spherical granules and larger, membrane-associated condensates as expected (Elaswad et al. 2024) (Figure 2A). In the primary screen, we initially identified six positive candidates among the sixteen genes: *let-711, copb-2*, *sar-1*, *sec-24.1*, *rpn-6.1*, and *pbs-7* (Table 1, Figure 2A). We detected one additional candidate gene, *rpl-3*, after further testing for false negatives with more stringent RNAi conditions. All seven genes were validated as positive candidates during multiple experiments performed by independent researchers. We qualitatively sorted the MEX-3 phenotypes into two categories: many ectopic spherical granules and increased membrane enrichment with a small number of ectopic granules (Table 1). We used the skewness tool in ImageJ as a proxy to measure the extent of MEX-3 condensation (described in methods). Our quantitative analysis revealed statistically significant increases in MEX-3 condensation after depletion of six of the seven genes (Figure 2B).

**Figure 2.**
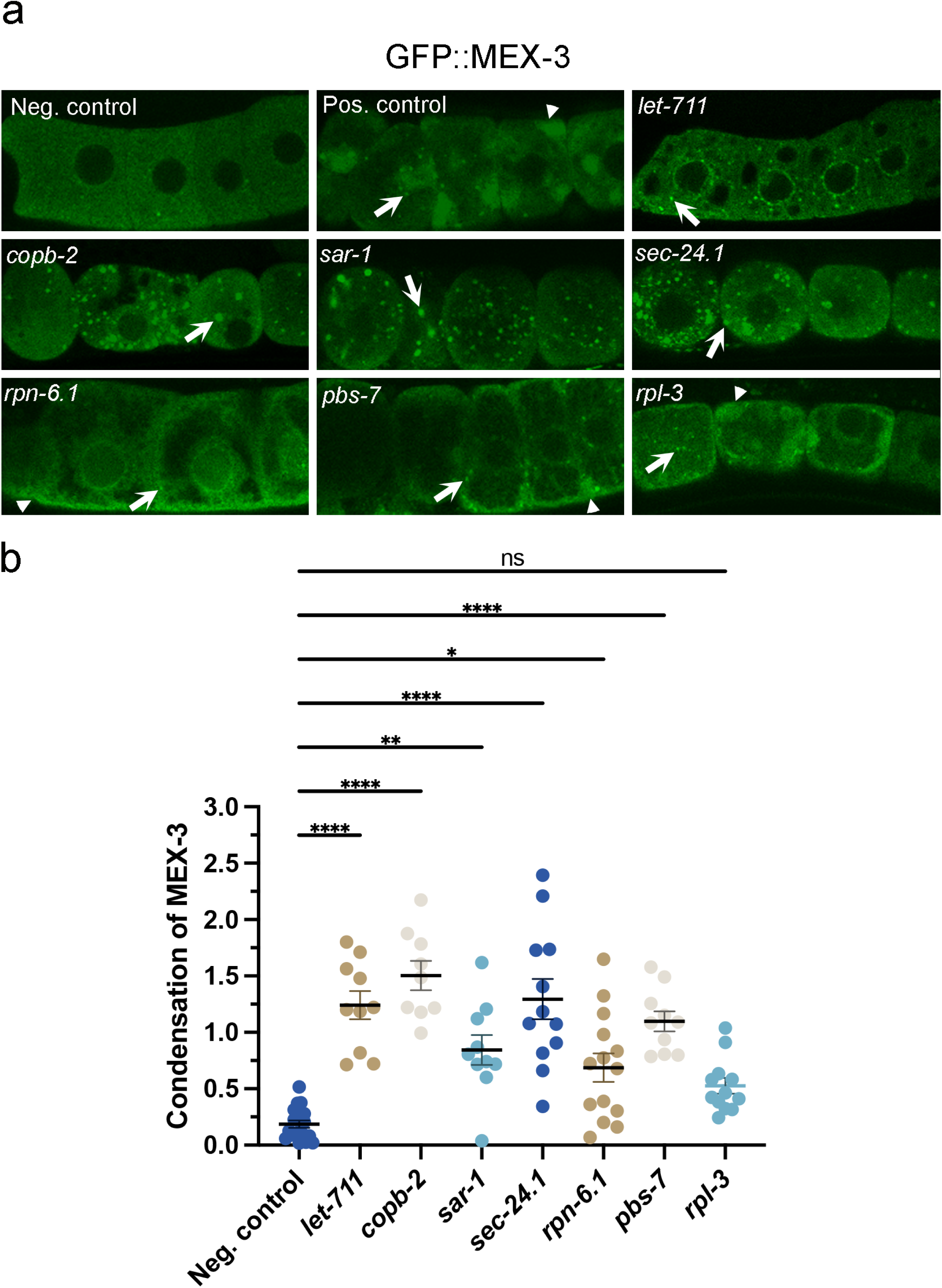
A subset of RHO-1 regulators prevents ectopic MEX-3 condensation in oocytes. A. Confocal images of GFP::MEX-3 in oocytes after RNAi depletions. MEX-3 is decondensed in the *lacZ(RNAi)* negative control. In the *cct-5(RNAi)* positive control, MEX-3 is condensed into small, spherical granules (arrows) and into larger heterogenous condensates enriched at membranes (arrowheads). Depletions of *let-711, copb-2, sar-1,* and *sec-24.1* resulted in ectopic spherical MEX-3 granules. Depletions of *rpn-6.1, pbs-7,* and *rpl-3* resulted in ectopic enrichments at membranes and small numbers of granules. All micrographs are single-Z plane confocal images, and the most proximal oocyte is oriented to the left. Scale bar is 10 µm. B. Significant increases in MEX-3 condensation was detected after all depletions except *rpl-3* using the Fiji Skewness tool. Each dot on the graph represents the average of the three skewness values for the −1 to −3 oocytes in each worm. Higher skewness values indicate less uniformity of GFP signal in an ROI which is a measure of increased condensation. Similar results were seen in both mid-focal and cortical Z slices (mid-focal values not shown). Kruskal-Wallis test. * P <0.05, ** P<0.01, **** P <0.0001. n=10-15.

### Novel regulators of MEX-3 phase transitions

#### Genes associated with protein degradation and protein synthesis

Depletion of three genes broadly involved in protein degradation and synthesis, *pbs-7, rpn-6.1,* and *rpl-3,* resulted in phenotypes similar to *cct-5*, i.e. increased membrane enrichment of MEX-3 with small, ectopic spherical granules (Figure 2). While the condensation of MEX-3 was not statistically increased after *rpl-3(RNAi)*, in half of the worms, condensation was increased above levels in the negative control. *rpl-3* encodes the ribosomal protein L3. Interestingly, the size of PGL-1 granules in embryos is positively regulated by ribosome genes (Updike and Strome 2009), which contrasts with the modest role we detected for the ribosome gene *rpl-3* inhibiting condensation of MEX-3 in oocytes. In different developmental contexts, active translation may modulate condensation or de-condensation of RNA-binding proteins.

The other two genes encode RPN-6.1 (ortholog of PSMD11), a proteasome 26S subunit involved in ubiquitin-dependent degradation, and PBS-7 (ortholog of PSMB4) which is part of the proteasome core complex. Ubiquitin binding has been implicated in modulating phase transitions of UBQLN2 *in vitro*, and 21 proteasomal and ubiquitin genes were identified in a screen to identify genes that promote condensation of the *C. elegans* P-granule protein PGL-1 in embryos (Updike and Strome 2009; Dao et al. 2018). Our results suggest the degradation machinery may also be needed to maintain a decondensed phase of MEX-3 in oocytes. If MEX-3 levels increase after these degradation gene depletions, the MEX-3 protein levels may accumulate beyond the concentration threshold required for phase separation, leading to ectopic condensation (Hyman et al. 2014; Alberti et al. 2017).

#### Genes associated with RNA processing

Prior to our study, the RNA helicase CGH-1/DDX6 was shown to prevent both ectopic MEX-3 condensates and ectopic RHO-1 aggregates in maturing oocytes (Langerak et al. 2019; Samaddar et al. 2021). In our primary screen, we tested two additional RHO-1 regulators that function broadly in RNA processing. We did not detect any changes in MEX-3 after depletion of *ess-2*; however, we detected a significant increase in ectopic MEX-3 condensates after depletion of *let-711/Not1*, an essential scaffolding protein in the CCR4-NOT deadenylase complex (Figure 2). This result suggested LET-711/NOT1 prevents ectopic condensation of MEX-3 in maturing oocytes. LET-711 has a similar role in modulating the phase transitions of two other RNA-binding proteins in *C. elegans*, preventing ectopic square, solid-like sheets of CAR-1/Lsm14 in arrested oocytes and inhibiting the condensation of PGL-1 germ granules in maturing oocytes (Updike and Strome 2009; Hubstenberger et al. 2015). Moreover, this role may be conserved; the yeast Not1p interacts with Dhh1p/DDX6 to promote disassembly of P-bodies, possibly through activation of the ATPase activity of Dhh1p (Sachdev et al. 2019). One possibility is that the disrupted deadenylation of mRNAs in oocytes may lead to an accumulation of undegraded maternal mRNA which may in turn, nucleate ectopic condensates of oogenic RNA-binding proteins.

#### Genes associated with vesicle-mediated trafficking

We identified three genes involved in vesicle-mediated trafficking between the ER and Golgi as novel inhibitors of ectopic MEX-3 condensation. After depletion of *copb-2, sar-1,* and *sec-24.1,* numerous spherical MEX-3 granules were detected in proximal oocytes that were unusually rounded (Figure 2). COPB-2/β’Cop is a subunit of the B subcomplex of the COPI-coat protein complex which functions to transport proteins within Golgi membranes and from the Golgi to the ER (Beck et al. 2009). While no reports to our knowledge have identified proteins in the COPI complex as modulators of RNA-binding protein phase transitions in oocytes, two COPI proteins, Sec27p/β’Cop and Sec 21p/γCop, limit the formation of P-bodies in yeast (Kilchert et al. 2010). Since COPI complexes can move improperly folded proteins from the Golgi to the ER, and various environmental stresses induce condensation of MEX-3, it is possible that deletion of *copb-2* in oocytes induces the Unfolded Protein Response or ER cellular stress response (Jud et al. 2008; Elaswad, Watkins, et al. 2022). An alternative possibility comes from yeast studies that support osmotic stress as a mediator of P-body induction in Sec mutants (Kilchert et al. 2010). Future studies can determine if COPB-2 functions to modulate RNA-binding protein phase transitions independently, or as part of, the activity of COPI vesicles, and the mechanism by which it acts.

SAR-1/Sar1 is an ER-exit-site small GTPase that is activated by the GEF SEC-12 to initiate COPII vesicle formation and enable transport of cargo from the ER to Golgi as part of the secretory pathway (Barlowe and Schekman 1993; Barlowe et al. 1993). SEC-24.1/Sec24D is recruited by SAR-1, along with SEC-23, to form the inner coat of the COPII vesicle, and SEC-24 recruits cargo into COPII vesicles (Matsuoka et al. 1998; Bi et al. 2002; Sato and Nakano 2007). When we first identified these two candidate regulators, we were not aware of published studies identifying proteins in the COPII complex as modulators of RBP phase transitions during oogenesis. However, four of the five proteins in the COPII complex were recently shown to regulate the organization of P-bodies in nurse cells which serve to support oocyte growth in *Drosophila* (Milano et al. 2025). After depleting expression of Sec23, Sec13, or Sec31, the condensate size for two P-body proteins increases; however, after depletion of Sar1, P-body proteins are less condensed in *Drosophila* nurse cells. Interestingly, our result where SAR-1 prevents condensation of MEX-3 in *C. elegans* oocytes contrasts with the *Drosophila* nurse cell result but is similar to the role of Sar1p in yeast where Sar1p limits the number of P-bodies in the cell (Kilchert et al. 2010). Future studies will determine whether these apparent differences underlie Sar1 functions that are independent of COPII vesicle trafficking, specific requirements in oocytes that differ from nurse cells, or simply differences in regulation among proteins.

### Regulatory network of RNA-binding protein phase transitions differs from protein aggregates

Among the nine RHO-1 regulators that were negatives in our primary screen were two genes involved in lysosome acidification (*vha-2* and *vha-12*), an ATP synthase in mitochondria that is required to decrease the membrane potential (*atp-3*), two calcium ion transport genes (*sca-1*, *itr-1*), two genes in the ESCRT complex (*vps-37, vps-54),* and an ER homeostasis gene (*spcs-1*). We selected one or two genes in each of these five GO categories in subsequent testing for false negatives. We performed RNAi in the GFP::MEX-3 and GFP::RHO-1 strains in parallel, with the RHO-1 strain acting as a positive control for depletion of gene expression since RHO-1 aggregates are expected (Samaddar et al. 2021).

In testing for false negatives, RNAi of *vha-12, vha-2, atp-3, sca-1,* and *vps-37* in GFP::RHO-1 worms resulted in RHO-1 aggregates in the proximal oocytes, suggesting at least modestly effective depletion of gene expression during RNAi (Figure 3A). However, after depleting each of these five genes, MEX-3 protein appeared unchanged from a largely decondensed state, similar to the negative control (Figure 3A). Our quantitative analysis did not reveal any significant increases in MEX-3 condensation (Figure 3B). In a minority of *vha-12(RNAi)* and *vha-2(RNAi)* worms (3 of 14 and 2 of 10, respectively), we detected a slight increase in the extent of MEX-3 condensation; however, some of the oocytes also appeared more stacked which may indicate a lack of sperm signal, a developmental context when MEX-3 condensation is known to occur (Jud et al. 2008) (Figure 1A). Alternatively, the low-penetrance, subtle phenotype may reflect a modest role for these two genes in preventing ectopic MEX-3 condensation in maturing oocytes or the limitations of depleting gene expression by RNAi. Taken together, our results suggest these five genes do not have major roles in preventing ectopic MEX-3 condensation. After RNAi of *spcs-1,* MEX-3 appeared largely decondensed as in the control (Table 1). Unfortunately, after multiple trials of *spcs-1(RNAi)* in GFP::RHO-1 worms, we did not detect the expected phenotype of ectopic aggregates (SFig. 1) (Samaddar et al. 2021). Therefore, we cannot make any conclusions regarding the possible regulation of MEX-3 phase transitions by *spcs-1*. Overall, the RHO-1 regulator genes required for V-ATPase assembly and docking, lysosomal acidification, regulation of mitochondrial membrane potential, and the ESCRT proteins, do not appear to have large roles in regulating phase transitions of MEX-3 in oocytes. Thus, we conclude the regulatory pathways in oocytes to prevent ectopic MEX-3 condensation and to inhibit ectopic RHO-1 aggregates are not completely coordinated.

**Figure 3.**
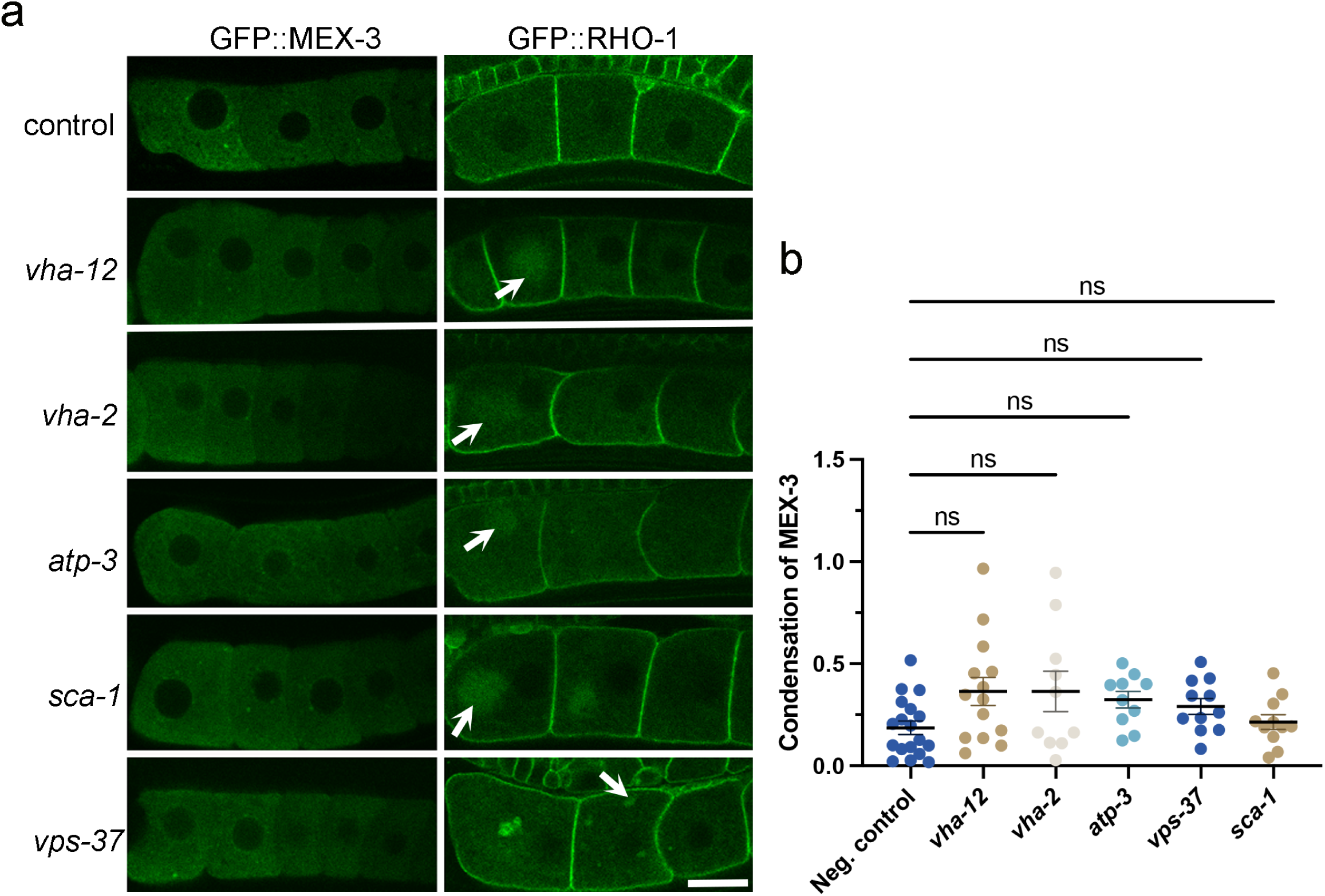
A subset of RHO-1 regulators is not required to prevent ectopic MEX-3 condensation in oocytes. A. *left column)* GFP::MEX-3 appears mostly decondensed in *lacZ(RNAi)* negative control oocytes and after depletion of five RHO-1 regulator genes. *right column)* The efficacy of each RNAi experiment was validated by parallel depletions in the GFP::RHO-1 strain. RHO-1 is enriched at the membrane of *lacZ(RNAi)* control oocytes. After depletion of the five genes, ectopic RHO-1 aggregates were detected (arrows). All micrographs are single-Z plane confocal images, and the most proximal oocyte is oriented to the left. Scale bar is 10 µm. B. No significant increases in MEX-3 condensation were detected after RNAi depletions using the FIJI Skewness tool. Each dot on the graph represents the average of the three skewness values for the −1 to −3 oocytes in each worm. Higher skewness values indicate less uniformity of GFP signal in an ROI which is a measure of increased condensation. Similar results were seen in both mid-focal and cortical Z slices (mid-focal values not shown). Kruskal-Wallis test. ns is not significant. n=10-15 worms.

### Conclusions and Considerations

High-quality oocytes are paramount to successful reproduction. Indeed, the ability of oocytes across species to maintain their quality during extended delays in fertilization is quite remarkable. Recent attention has focused on identifying mechanisms that protect oocytes from cytosolic damage such as protein misfolding and aggregation (Bohnert and Kenyon 2017; Samaddar et al. 2021; Harasimov et al. 2024; Rosswag de Souza et al. 2025). At the same time, we have long known the requirement in oocytes to protect maternal mRNAs and their partner RNA-binding proteins. However, the extent to which the temporally linked processes of clearance of damaged proteins and maintenance of RNP complexes are coordinated at a molecular level has not been examined. In this targeted genetic screen, we identified six novel regulators of MEX-3 phase transitions and obtained strong evidence that the regulatory network of protein aggregate clearance is distinct from the regulation of MEX-3 phase transitions in the oocyte.

In addition to the discussion of our results above, we noted an intriguing correlation between genes required to inhibit ectopic MEX-3 condensation and for normal ER architecture. Our prior studies describe three inhibitors of MEX-3 condensation, the CCT chaperonin, actin, and CGH-1, that are also required for normal ER morphology in oocytes (Langerak et al. 2019; Elaswad et al. 2024). Here, we add *copb-2* and *pbs-7* to the list of genes that modulate both ER morphology and MEX-3 condensation (Samaddar et al. 2021). Conversely, genes that appear to prevent RHO-1 aggregation but not MEX-3 condensation, *vha-12, vha-2,* and *atp-3,* do not affect ER morphology. These findings are consistent with a model where expanded ER sheets in oocytes contribute to MEX-3 condensation by restricting molecular diffusion to a two-dimensional plane, thereby lowering the concentration threshold for protein condensation (Snead and Gladfelter 2019; Elaswad et al. 2024).

Our identification of novel inhibitors of ectopic MEX-3 condensation advances our understanding of RNA-binding protein phase transitions in maturing oocytes. The results provide a foundation for future studies to probe the mechanisms by which these genes modulate MEX-3 phase and determine whether any genes have more global roles in regulating RNA-binding proteins phase transitions in maturing oocytes. Because aberrant phase transitions of hMex3 alter its interactions with P-bodies that modulate mRNA degradation and are associated with the progression of colorectal cancer (Chen et al. 2024), our findings may have applications beyond oocyte quality and fertility.

**SFigure 1.** RNAi depletion of *spcs-1* did not reveal phenotypes for MEX-3 or RHO-1. Confocal micrographs of negative control, *lacZ(RNAi),* and *spcs-1(RNAi)* in GFP::MEX-3 and GFP::RHO-1 strains. No RHO-1 aggregates were detected, indicating the RNAi may not have been effective.

## ACKNOWLEDGEMENTS

We thank Dr. David Greenstein and the *Caenorhabditis* Genetics Center, which is funded by NIH Office of Research Infrastructure Programs (P40 OD010440), for strains. We thank Drs. Xantha Karp and Lisa Petrella for helpful discussion. We acknowledge the CMU College of Science and Engineering for support of graduate student M.T.E. This work was funded by NIH grant 1R15GM147844-01 to J.A.S.

## Literature Cited

Alberti S, Mateju D, Mediani L, Carra S. 2017. Granulostasis: Protein quality control of RNP granules. Front Mol Neurosci. 10. 10.3389/fnmol.2017.00084

Audhya A, Hyndman F, McLeod IX, Maddox AS, Yates JR 3rd, Desai A, Oegema K. 2005. A complex containing the Sm protein CAR-1 and the RNA helicase CGH-1 is required for embryonic cytokinesis in Caenorhabditis elegans. Journal of Cell Biology. 171(2):267–279. 10.1083/jcb.200506124

Barlowe C, Schekman R. 1993. SEC12 encodes a guanine-nucleotide-exchange factor essential for transport vesicle budding from the ER. Nature. 365:347–349

Barlowe C, d’Enfert C, Schekman R. 1993. Purification and characterization of SAR1p, a small GTP-binding protein required for transport vesicle formation from the endoplasmic reticulum. J Biol Chem. 268(2):873–9.

Bi X, Corpina RA, Goldberg J. 2002. Structure of the Sec23/24-Sar1 pre-budding complex of the COPII vesicle coat. Nature. 419(6904):271–7. doi: 10.1038/nature01040.

Beck R, Ravet M, Wieland FT, Cassel D. 2009. The COPI system: Molecular mechanisms and function. FEBS Lett. 583(17):2701–2709. 10.1016/j.febslet.2009.07.032

Boag PR, Nakamura A, Blackwell TK. 2005. A conserved RNA-protein complex component involved in physiological germline apoptosis regulation in C. elegans. Development. 132(22):4975–4986. 10.1242/dev.02060

Bohnert AK, Kenyon C. 2017. A lysosomal switch triggers proteostasis renewal in the immortal C. Elegans germ lineage. Nature. 551(7682):629–633. 10.1038/nature24620

Bose M, Lampe M, Mahamid J, Ephrussi A. 2022. Liquid-to-solid phase transition of oskar ribonucleoprotein granules is essential for their function in Drosophila embryonic development. Cell. 185(8):1308–1324.e23. 10.1016/j.cell.2022.02.022

Brenner S. 1974. The genetics of Caenorhabditis elegans. Genetics 77(1):71–94. 10.1093/genetics/77.1.71.

Buchet-Poyau K, Courchet J, Le Hir H, Séraphin B, Scoazec JY, Duret L, Domon-Dell C, Freund JN, Billaud M. 2007. Identification and characterization of human Mex-3 proteins, a novel family of evolutionarily conserved RNA-binding proteins differentially localized to processing bodies. Nucleic Acids Res. 35(4):1289–1300. 10.1093/nar/gkm016

Chen RX, Xu SD, Deng MH, Hao SH, Chen JW, Ma XD, Zhuang WT, Cao JH, Lv YR, Lin JL, et al. 2024. Mex-3 RNA binding family member A (MEX3A)/circMPP6 complex promotes colorectal cancer progression by inhibiting autophagy. Signal Transduct Target Ther. 9(1). 10.1038/s41392-024-01787-3

Conti M, Kunitomi C. 2024. A genome-wide perspective of the maternal mRNA translation program during oocyte development. Semin Cell Dev Biol. 154:88–98. 10.1016/j.semcdb.2023.03.003

Cheng S, Altmeppen G, So C, Welp LM, Penir S, Ruhwedel T, Menelaou K, Harasimov K, Stützer A, Blayney M, Elder K, Möbius W, Urlaub H, Schuh M. 2022. Mammalian oocytes store mRNAs in a mitochondria-associated membraneless compartment. Science (1979). 378(6617). 10.1126/science.abq4835

Dao TP, Kolaitis RM, Kim HJ, O’Donovan K, Martyniak B, Colicino E, Hehnly H, Taylor JP, Castañeda CA. 2018. Ubiquitin Modulates Liquid-Liquid Phase Separation of UBQLN2 via Disruption of Multivalent Interactions. Mol Cell. 69(6):965–978.e6. 10.1016/j.molcel.2018.02.004

Donnini M, Lapucci A, Papucci L, Witort E, Jacquier A, Brewer G, Nicolin A, Capaccioli S, Schiavone N. 2004. Identification of TINO: A new evolutionarily conserved BCL-2 AU-rich element RNA-binding protein. Journal of Biological Chemistry. 279(19):20154–20166. 10.1074/jbc.M314071200

Draper BW, Mello CC, Bowerman B, Hardin J, Priess JR. 1996. MEX-3 Is a KH Domain Protein That Regulates Blastomere Identity in Early C. elegans Embryos. Cell.; 87(2):205–216. doi:10.1016/s0092-8674(00)81339-2

Elaswad MT, Watkins BM, Sharp KG, Munderloh C, Schisa JA. 2022. Large RNP granules in Caenorhabditis elegans oocytes have distinct phases of RNA-binding proteins. G3: Genes, Genomes, Genetics. 12(9). 10.1093/g3journal/jkac173

Elaswad MT, Munderloh C, Watkins BM, Sharp KG, Breton E, Schisa JA. 2022. Imaging-Associated stress causes divergent phase transitions of RNA-binding proteins in the Caenorhabditis elegans germ line. G3: Genes, Genomes, Genetics. 12(9). 10.1093/g3journal/jkac172

Elaswad MT, Gao M, Tice VE, Bright CG, Thomas GM, Munderloh C, Trombley NJ, Haddad CN, Johnson UG, Cichon AN, Schisa JA. 2024. The CCT chaperonin and actin modulate the ER and RNA-binding protein condensation during oogenesis and maintain translational repression of maternal mRNA and oocyte quality. Mol Biol Cell. 35(10). 10.1091/mbc.E24-05-0216

Flemr M, Ma J, Schultz RM, Svoboda P. 2010. P-body loss is concomitant with formation of a messenger RNA storage domain in mouse oocytes. Biol Reprod. 82(5):1008–1017. 10.1095/biolreprod.109.082057

Holehouse AS, Alberti S. 2025. Molecular determinants of condensate composition. Mol Cell. 85(2):290–308. 10.1016/j.molcel.2024.12.021

Hubstenberger A, Noble SL, Cameron C, Evans TC. 2013. Translation repressors, an RNA helicase, and developmental cues control RNP phase transitions during early development. Dev Cell. 27(2):161–173. 10.1016/j.devcel.2013.09.024

Hubstenberger A, Cameron C, Noble SL, Keenan S, Evans TC. 2015. Modifiers of solid RNP granules control normal RNP dynamics and mRNA activity in early development. Journal of Cell Biology. 211(3):703–716. 10.1083/jcb.201504044

Hyman AA, Weber CA, Jülicher F. 2014. Liquid-liquid phase separation in biology. Annu Rev Cell Dev Biol. 30:39–58. 10.1146/annurev-cellbio-100913-013325

Jud M, Razelun J, Bickel J, Czerwinski M, Schisa JA. 2007. Conservation of large foci formation in arrested oocytes of Caenorhabditis nematodes. Dev Genes Evol. 217(3):221–226. 10.1007/s00427-006-0130-3

Jud MC, Czerwinski MJ, Wood MP, Young RA, Gallo CM, Bickel JS, Petty EL, Mason JM, Little BA, Padilla PA, Schisa JA. 2008. Large P body-like RNPs form in C. elegans oocytes in response to arrested ovulation, heat shock, osmotic stress, and anoxia and are regulated by the major sperm protein pathway. Dev Biol. 318(1):38–51. 10.1016/j.ydbio.2008.02.059

Kamath RS, Ahringer J. 2003. Genome-wide RNAi screening in Caenorhabditis elegans. Methods. 30(4):313–321. 10.1016/S1046-2023(03)00050-1

Kilchert C, Weidner J, Prescianotto-Baschong C, Spang A. 2010. Defects in the Secretory Pathway and High Ca 2 Induce Multiple P-bodies. Mol Biol Cell. 21:2624–2638 10.1091/mbc.E10

Langerak S, Trombley A, Patterson JR, Leroux D, Couch A, Wood MP, Schisa JA. 2019. Remodeling of the endoplasmic reticulum in Caenorhabditis elegans oocytes is regulated by CGH-1. Genesis. 57(2):e23267 https://www.ncbi.nlm.nih.gov/pubmed/30489010. 10.1002/dvg.23267

Ladomery M, Wade E, Sommerville J. 1997. Xp54, the Xenopus homologue of human RNA helicase p54, is an integral component of stored mRNP particles in oocytes. Nucleic Acids Res. 1;25(5):965–73. doi: 10.1093/nar/25.5.965.

Matsuoka K, Morimitsu Y, Uchida K, Schekman R. 1998. Coat Assembly Directs v-SNARE Concentration into Synthetic COPII Vesicles. Molec Cell 2(5) 703–708. 10.1016/S1097-2765(00)80168-9.

Milano SN, Bayer LV, Ko JJ, Casella CE, Bratu DP. 2025. The role of ER exit sites in maintaining P-body organization and integrity during Drosophila melanogaster oogenesis. EMBO Rep. 26(2):494–520. 10.1038/s44319-024-00344-x

Nakamura A, Amikura R, Hanyu K, Kobayashi S. 2001. Me31B silences translation of oocyte-localizing RNAs through the formation of cytoplasmic RNP complex during Drosophila oogenesis. Development. 128:3233–3242. doi: 10.1242/dev.128.17.3233.

Navarro RE, Shim EY, Kohara Y, Singson A, Blackwell TK. 2001. cgh-1, a conserved predicted RNA helicase required for gametogenesis and protection from physiological germline apoptosis in C. elegans. Development. 128(17):3221–32. doi: 10.1242/dev.128.17.3221.2001.

Noble SL, Allen BL, Goh LK, Nordick K, Evans TC. 2008. Maternal mRNAs are regulated by diverse P body-related mRNP granules during early Caenorhabditis elegans development. Journal of Cell Biology. 182(3):559–572. 10.1083/jcb.200802128

Patterson JR, Wood MP, Schisa JA. 2011. Assembly of RNP granules in stressed and aging oocytes requires nucleoporins and is coordinated with nuclear membrane blebbing. Dev Biol. 353(2):173–185. 10.1016/j.ydbio.2011.02.028

Rosswag de Souza, Boke E, Zaffagnini G. 2025. Proteostasis in cellular dormancy: lessons from yeast to oocytes. Trends Biochem Sci 50(8) 646–662. 10.1016/j.tibs.2025.05.004

Sachdev R, Hondele M, Linsenmeier M, Vallotton P, Mugler CF, Arosio P, Weis K. 2019. Pat1 promotes processing body assembly by enhancing the phase separation of the DEAD-box ATPase Dhh1 and RNA. Elife. e41415 https://elifesciences.org/articles/41415#abstract. 10.7554/eLife.41415.001

Samaddar M, Goudeau J, Sanchez M, Hall DH, Bohnert KA, Ingaramo M, Kenyon C. 2021. A genetic screen identifies new steps in oocyte maturation that enhance proteostasis in the immortal germ lineage. Elife. 10. 10.7554/ELIFE.62653

Sato K, Nakano A. 2007. Mechanisms of COPII vesicle formation and protein sorting. FEBS Lett. 581(11):2076–2082. 10.1016/j.febslet.2007.01.091

Schisa JA, Pitt JN, Priess JR. 2001. Analysis of RNA associated with P granules in germ cells of C. elegans adults. Development 128:1287–1298. 10.1242/dev.128.8.1287

Snead WT, Gladfelter AS. 2019. The Control Centers of Biomolecular Phase Separation: How Membrane Surfaces, PTMs, and Active Processes Regulate Condensation. Mol Cell. 76(2):295–305. 10.1016/j.molcel.2019.09.016

Spike CA, Coetzee D, Eichten C, Wang X, Hansen D, Greenstein D. 2014. The TRIM-NHL protein LIN-41 and the OMA RNA-binding proteins antagonistically control the prophase-to-metaphase transition and growth of Caenorhabditis elegans oocytes. Genetics. 198(4):1535–1558. 10.1534/genetics.114.168831

Squirrell JM, Eggers ZT, Luedke N, Saari B, Grimson A, Lyons GE, Anderson P, White JG. 2006. CAR-1, a Protein That Localizes with the mRNA Decapping Component DCAP-1, Is Required for Cytokinesis and ER Organization in Caenorhabditis elegans Embryos □ D □ V. Mol Biol Cell. 17:336–344. 10.1091/mbc.E05-09

Timmons L, Fire A. 1998. Specific interference by ingested dsRNA. Nature 395(6705):854. doi:10.1038/27579.

Tsukamoto T, Gearhart MD, Spike CA, Huelgas-Morales G, Mews M, Boag PR, Beilharz TH, Greenstein D. 2017. LIN-41 and OMA Ribonucleoprotein Complexes Mediate a Translational Repression-to-Activation Switch Controlling Oocyte Meiotic Maturation and the Oocyte-to-Embryo Transition in *Caenorhabditis elegans*. Genetics. 206(4):2007–2039. 10.1534/genetics.117.203174

Updike DL, Strome S. 2009. A genomewide RNAi screen for genes that affect the stability, distribution and function of P granules in Caenorhabditis elegans. Genetics. 183(4):1397–1419. 10.1534/genetics.109.110171

Wood MP, Hollis A, Severance AL, Karrick ML, Schisa JA. 2016. RNAi screen identifies novel regulators of RNP granules in the Caenorhabditis elegans germ line. G3: Genes, Genomes, Genetics. 6(8):2643–2654. 10.1534/g3.116.031559

